# Identification of a novel gammaherpesvirus in Canada lynx (*Lynx canadensis*)

**DOI:** 10.1101/579607

**Authors:** Liam D. Hendrikse, Ankita Kambli, Caroline Kayko, Marta Canuti, Bruce Rodrigues, Brian Stevens, Jennifer Vashon, Andrew S. Lang, David B. Needle, Ryan M. Troyer

## Abstract

Gammaherpesviruses (GHVs) infect many animal species and are associated with lymphoproliferative disorders in some. Previously, we identified several novel GHVs in North American felids, however a GHV had never been identified in Canada lynx (*Lynx canadensis*). We therefore hypothesized the existence of an unidentified GHV in lynx. Using degenerate nested and subsequently virus-specific PCR, we amplified and sequenced 3.4 kb of DNA from a novel GHV in lynx, which we named *Lynx canadensis* gammaherpesvirus 1 (LcaGHV1). We then estimated the prevalence of LcaGHV1 in lynx by developing a PCR-based assay and detected LcaGHV1 DNA in 36% (95% CI: 22-53%) of lynx spleen DNA samples from Maine, USA and 17% (95% CI: 8-31%) from Newfoundland, Canada. Phylogenetic analysis determined that LcaGHV1 is a distinct GHV species belonging to the genus *Percavirus*. The potential ecologic and pathologic consequences of this novel virus for Canada lynx populations warrant further study.

## Introduction

Gammaherpesviruses (GHVs) belong to the *Herpesviridae* family (subfamily *Gammaherpesvirinae*), a large group of double-stranded DNA viruses (King et al., 2012). GHVs can be further divided into four genera: *Lymphocryptovirus, Macavirus, Rhadinovirus*, and *Percavirus* (King et al., 2012). GHVs are a widely distributed group of viruses; they have been discovered in a plethora of mammals and even in invertebrates (Barton et al., 2011; Cabello et al., 2013; Chmielewicz et al., 2003, 2001, Ehlers et al., 2008, 2007; McGeoch et al., 2006; Shabman et al., 2016; Wibbelt et al., 2007; Wu et al., 2012). The evolution of GHVs is thought to have been tied to that of their respective hosts, and thus much of the diversity of GHVs may be explained by host speciation events (Jung and Speck, 2013; Wertheim et al., 2014). As a result, GHVs are well adapted to their specific host and diseases are often mild. The most well characterized diseases caused by GHVs are lymphocyte proliferative disorders (Ackermann, 2006). In humans, lymphocyte proliferative disorders are caused by Epstein-Barr virus (EBV) and Kaposi’s sarcoma-associated herpesvirus (KHSV), which are members of the *Lymphocryptovirus* and *Rhadinovirus* genera, respectively. Diseases associated with these viruses occur primarily in immunocompromised individuals (Cesarman, 2011; Mesri et al., 2010). Macaviruses such as alcelaphine gammaherpesvirus 1 (AlGHV1) and ovine gammaherpesvirus 2 (OvGHV2) can cause malignant catarrhal fever, a virulent lymphoproliferative disease, when transmitted to ungulates that lack immunity to these agents (Russell et al., 2009). The percaviruses equine gammaherpesvirus 2 (EGHV2) and equine gammaherpesvirus 5 (EGHV5) have been associated with respiratory disease in their natural horse hosts (Fortier et al., 2010), but little is known about the disease-causing potential of other percaviruses. Regardless of genus, infection by a GHV typically results in a short acute phase followed by an extended latency phase (Barton et al., 2011). As with all herpesviruses, GHVs use latency to evade immune system detection and as such, GHV infections are life-long (Ackermann, 2006; Barton et al., 2011). GHVs typically establish latency in lymphocytes, and they have been found in many tissues including the spleen, small intestine, lung, bone marrow, and more (Ackermann, 2006; Barton et al., 2011; Beatty et al., 2014; Hughes et al., 2010; Jung and Speck, 2013; Speck and Ganem, 2010). While contributing to a lifelong infection, this characteristic also assists in the discovery of GHVs, since their genomes can be found in tissues isolated from infected individuals, symptomatic or not.

We have previously described the first domestic cat GHV; *Felis catus* gammaherpesvirus 1 (FcaGHV1), which has since been found in cats from North America, South America, Europe, Australia, and Asia (Beatty et al., 2014; Ertl et al., 2015; Kurissio et al., 2018; Tateno et al., 2017; Troyer et al., 2014). We also identified novel GHV species in bobcats (*Lynx rufus*, LruGHV1 and LruGHV2), pumas (*Puma concolor*, PcoGHV1), and ocelots (*Leopardus pardalis*, LpaGHV1) (Lozano et al., 2015; Troyer et al., 2014). In addition, a previous study identified a GHV in an African lion (*Panthera leo*, PleoGHV1) (Ehlers et al., 2008). All of the previously identified feline GHVs have been found to cluster within the genus *Percavirus*, with the exceptions of PcoGHV1 and PleoGHV1, which belong to the genus *Rhadinovirus* (Lozano et al., 2015; Troyer et al., 2014). There has never been a GHV identified in Canada lynx (*Lynx canadensis*), referred to subsequently in this report as “lynx”. Lynx can be found across Canada and the northern United States. On the island of Newfoundland, the lynx population is isolated enough to form a lynx subspecies, *Lynx canadensis subsolanus* (Canuti et al., 2017; Koen et al., 2015; Row et al., 2012; Van Zyll De Jong, 1975). This isolation has made the Newfoundland lynx population particularly interesting for the study of pathogen evolution and epidemiology (Canuti et al., 2017; Koen et al., 2015; Van Zyll De Jong, 1975). The presence of at least one GHV in all of the other feline species tested to date provides support for the existence of an unidentified GHV in lynx. We thus hypothesized that there may be an unidentified lynx GHV, and further, that we would be able to detect this virus using PCR-based techniques on DNA from lymphocyte-containing lynx tissues.

In this study, we utilized several degenerate pan-GHV PCR protocols to screen lynx for novel viruses. We identified a previously unknown GHV glycoprotein B (gB) sequence and subsequently identified an unknown DNA polymerase (DNApol) gene as well. Using virus-specific primers, we amplified and sequenced a 3.4 kb region of this putative virus’ genome and conducted phylogenetic analysis to determine its relationship to other viruses. We then estimated the prevalence of this virus in the lynx populations of Maine and Newfoundland using a virus-specific PCR-based assay developed in this study. Finally, we utilized the prevalence data to statistically determine risk factors for infection.

## Methods

### Sample collection and DNA preparation

*Lynx canadensis* found dead or accidentally trapped in Maine, USA were collected over a 2-year period from 2016 to 2017 by the Maine Department of Inland Fisheries and Wildlife. The appropriate permits were obtained prior to sample collection and sex and location information were recorded when they could be reliably determined. Gross and histologic pathology was performed on all animals by the New Hampshire Veterinary Diagnostic Laboratory. *Lynx canadensis subsolanus* spleen samples were collected over a 5-year period from 2012 to 2016 from Newfoundland island, Canada as part of other ongoing studies (Canuti et al., 2017). Receipt of the lynx spleen samples was covered under protocols 14-04-AL and 15-04-AL (“Samples from wild fur-bearing mammals”) from the Memorial University Institutional Animal Care Committee. The sex and location of these animals within Newfoundland could not be reliably determined. All tissue samples were stored at −80°C prior to DNA extraction. Total DNA was extracted from thawed tissue samples using a DNeasy blood and tissue kit (Qiagen, Toronto, Ontario, Canada). Stringent measures were taken to avoid sample cross-contamination, including use of separate sterile instruments and plastics for disruption of each tissue sample.

### GAPDH PCR

The presence of intact, amplifiable, and inhibitor-free template DNA in each sample was confirmed using PCR for carnivore glyceraldehyde-3-phosphate dehydrogenase (GAPDH). Reactions of 20 μL total were prepared, containing 10 μL GoTaq Hot Start Green Master Mix (Promega, Madison, Wisconsin, USA), 0.4 μM primers GAPcons-F and GAPcons-R, and 1 μL of template DNA (see Table 1 for primer sequences). Cycling conditions were as follows: initial denaturing at 94°C for 3 min, followed by 35 cycles of 94°C for 30 s, 55°C for 30 s, and 72°C for 30 s and then a 10-minute extension at 72°C. PCR products of expected size (166 bp) were visualized by electrophoresis on a 1.5% agarose gel. Water control reactions were negative.

**Table 1.**
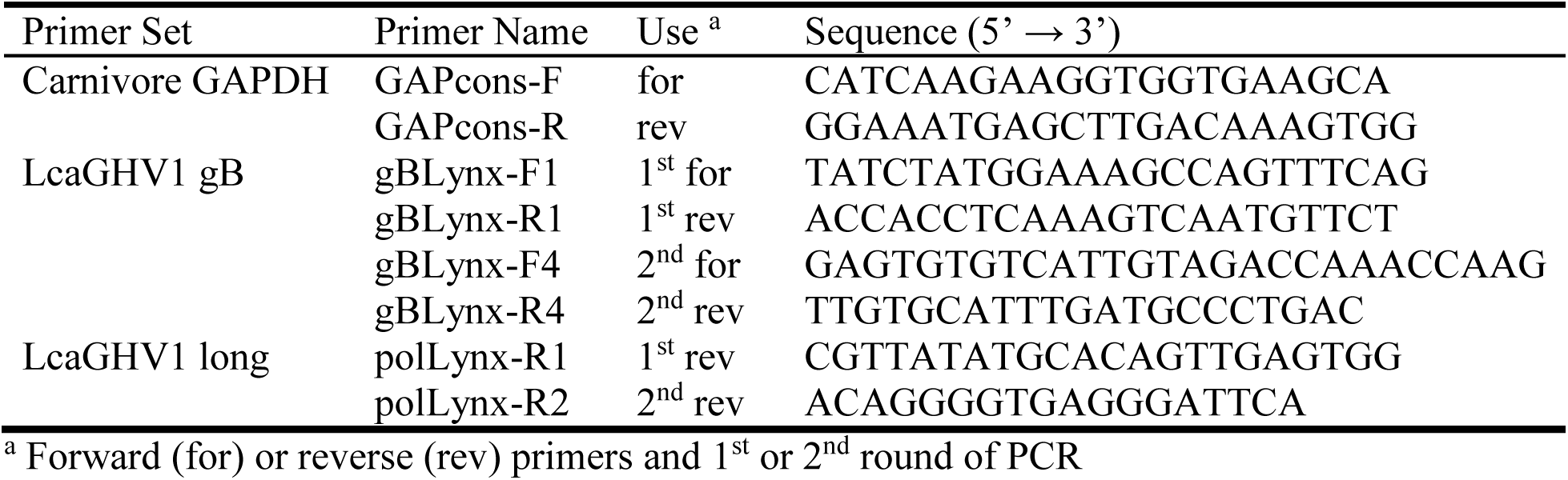
Primers designed in this study

### Amplification of GHV glycoprotein B (gB) sequence using degenerate primers

The presence of GHV DNA was determined using degenerate nested pan-GHV PCR to amplify a portion of the gB gene using the previously described primer set RH-gB (Ehlers et al., 2008). In the first round, 10 μL template DNA (100 to 1000 ng total) was added to 50 μL reaction mixtures containing 2 units Platinum Taq polymerase (ThermoFisher Scientific, Waltham, Massachusetts, USA), 1 μM primers 2759s and 2762as, 1.5 mM MgCl_2_, 0.2 mM deoxynucleoside triphosphates, and 1X PCR buffer (ThermoFisher Scientific). Cycling conditions were as follows: initial denaturing at 94°C for 2 min, followed by 45 cycles of 94°C for 30 s, 46°C for 30 s, and 72°C for 30 s and then a 7-minute extension at 72°C. In the second round, 2 μL of the first-round reaction product was used as the template and reactions were conducted under conditions identical to those described above using primers 2760s and 2761as. Stringent measures were taken to avoid cross-contamination, including use of separate areas for reaction mix preparation and template addition. The second-round products were visualized by agarose gel electrophoresis. Negative-control reactions (water template) were consistently negative. PCR products of the expected ∼500 bp were purified from the gel using a QIAquick Gel Extraction kit (Qiagen) and sequenced in both directions by Eurofins Genomics (Toronto, Ontario, Canada). After removal of the primer sequence, unique 452 bp gB sequences were compared to other known GHV gB sequences using BioEdit (Hall, 1999) and NCBI BLAST programs.

### Amplification of GHV DNA polymerase (DNApol) sequence using degenerate primers

The DNApol gene was amplified by degenerate nested PCR using previously described percavirus-specific first round primers LSSG and GDTD2 and second round primers VYGA2 and GDTD2 (Troyer et al., 2014). Amplification of DNApol was also attempted using pan-HV primers DFASA and GDTD1B in the first round and VYGA and GDTD1B in the second round (Rose et al., 1997). The reaction mix was identical to that of the gB PCR above, however 5 μL DNA (50 to 500 ng total) was added in the first round instead of 10 μL. Cycling conditions for both rounds were as follows: initial denaturing at 94°C for 2 min, followed by 45 cycles of 94°C for 60 s, 55°C (60°C for pan-HV primers) for 60 s, and 72°C for 60 s and then a 7-minute extension at 72°C. In the second round, 2 μL of the first-round reaction product was used as the template. Second-round products were visualized by agarose gel electrophoresis. PCR products of ∼236 bp were purified using a QIAquick Gel Extraction kit (Qiagen) and sequenced in both directions by Eurofins Genomics. After removal of the primer sequences, unique 172 bp DNApol sequences were compared to other known GHV DNApol sequences using the BioEdit (Hall, 1999) and NCBI BLAST programs.

### Long-distance PCR and sequencing

A sequence spanning from gB to DNApol (∼3.4 kb) was amplified by nested PCR (Figure 1A). In the first round, 5 μL DNA (125 to 500 ng total) was added to 50 μL reaction mixtures containing 2 units High Fidelity Platinum Taq polymerase (ThermoFisher Scientific), 0.4 μM primers gBLynx-F1 and polLynx-R1, 2 mM MgSO_4_, 0.2 mM deoxynucleoside triphosphates, and 1X PCR buffer (ThermoFisher Scientific). Cycling conditions were as follows: initial denaturing at 94°C for 2 min, followed by 40 cycles of 94°C for 30 s, 55°C for 30 s, and 68°C for 4 min and then a 7-minute extension at 68°C. In the second round, 2 μL of the first-round reaction product was used as the template and reactions were conducted under conditions identical to those described above using primers gBLynx-F4 and polLynx-R2. PCR products were purified using Agencourt AMPure XP reagent (Beckman Coulter, Brea, California, USA). Sequences from other known GHVs were aligned at the 3.4 kb region spanning from gB to DNApol gene. Using this alignment, sequencing primers were designed against regions of conserved nucleotide sequence. These primers were used to sequence the 3.4 kb PCR product in a series of overlapping reads performed by Eurofins Genomics. The reads obtained from sequencing were aligned using BioEdit (Hall, 1999), and assembled into a contiguous sequence.

**Figure 1.**
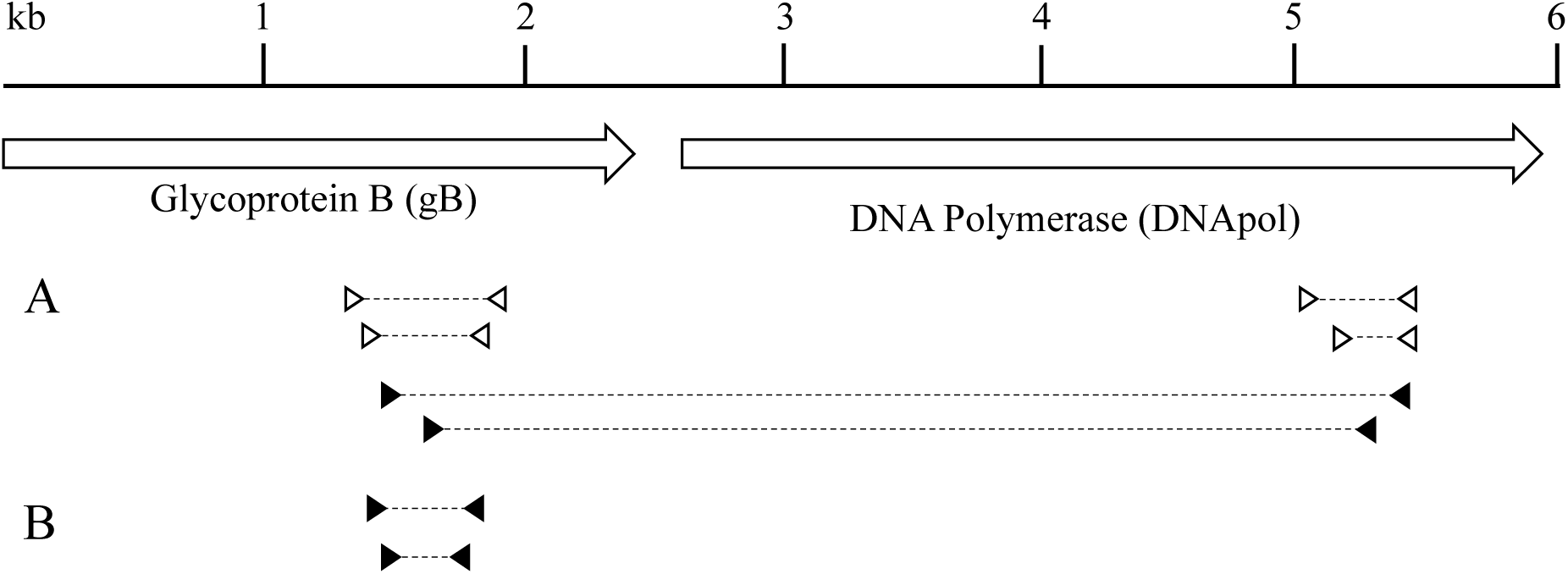
PCR and sequencing strategy. (A) Nested primer sets are indicated by triangles, with their respective PCR products indicated with a dashed line. Open triangles indicate degenerate primers that were not designed in this study, while closed triangles indicate virus-specific primers designed in this study. Degenerate primer sets were used for initial detection of novel GHVs and their sequences were used to design the virus-specific primers. Virus-specific primers were used to amplify an extended 3.4 kb portion of the viral genome. (B) LcaGHV1 screening primers. Nested primers were designed using the sequence obtained from the gB gene. (Figure adapted from Troyer et al., 2014)

### LcaGHV1 detection PCR assay

Based on the sequences obtained for the gB gene, LcaGHV1-specific primers were designed using Primer3 (Table 1). To detect LcaGHV1, nested PCR was performed (Figure 1B). In the first round, 5 μL DNA (50 to 500 ng total) was added to 50 μL reaction mixtures containing 2 units Platinum Taq polymerase (ThermoFisher Scientific), 0.4 μM primers gBLynx-F1 and gBLynx-R1, 1.5 mM MgCl_2_, 0.2 mM deoxynucleoside triphosphates, and 1X PCR buffer (ThermoFisher Scientific). Cycling conditions were as follows: initial denaturing at 94°C for 2 min, followed by 35 cycles of 94°C for 30 s, 58°C for 30 s, and 72°C for 30 s and then a 7 min extension at 72°C. In the second round, 2 μL of the first-round reaction product were used as the template for amplification with primers gBLynx-F4 and gBLynx-R4. The cycling conditions were identical to the first round, except an annealing temperature of 66°C was used. PCR products of 229 bp were purified using a QIAquick Gel Extraction kit (Qiagen) and sequenced by Eurofins Genomics. Unique sequences of ∼220 bp were aligned with the 452 bp LcaGHV1 gB gene sequence found previously in order to verify the identity of the virus.

### Statistical Analysis

We calculated the prevalence of infection by dividing the number of positive samples by the total number of samples tested and then utilized maximum likelihood estimation to determine confidence intervals (CIs) for prevalence. We examined potential risk factors for LcaGHV1 infection using generalized linear models with binomial error distributions.

### Phylogenetic Analysis

Phylogenetic analysis was performed on concatenated GHV gB and DNApol amino acid sequences. LcaGHV1 gB and DNApol amino acid sequences were aligned with sequences from 30 other HVs using the T-Coffee multiple alignment program with default parameters (Notredame et al., 2000). All positions with gaps and areas with weak support for the alignment were manually removed. The resulting gB and DNApol alignments were concatenated into a single alignment of 938 amino acids. The data were input into the DataMonkey (Delport et al., 2010a) server to estimate the best-fit model of amino acid substitution (Delport et al., 2010b). Maximum likelihood (ML) phylogenetic analysis was conducted using the PhyML program (Guindon and Gascuel, 2003) applied in Geneious Pro (version 11.1.2) software (Biomatters, Auckland, New Zealand) with Le and Gascuel (LG) (Le and Gascuel, 2008) substitution model and gamma distribution with five discrete categories. ML trees were constructed using a neighbour-joining starting tree followed by a heuristic search using the nearest-neighbour interchange algorithm. The betaherpesvirus human cytomegalovirus (HCMV; human herpesvirus 5 [HHV5]) was used as an outgroup to root the tree. Bootstrap analyses were performed with 100 iterations to evaluate the support for each node.

### Nucleotide Sequence Accession Numbers

The *Lynx canadensis* gammaherpesvirus 1 (LcaGHV1) nucleotide sequences found in this study were deposited into GenBank (http://www.ncbi.nlm.nih.gov/GenBank/index.html) under the following accession numbers: *Lynx canadensis* isolate, <u>MH520115</u>; and *Lynx canadensis subsolanus* isolate, <u>MH520116</u>.

## Results

### Detection of a novel gammaherpesvirus sequence in lynx tissues

To search for gammaherpesviruses (GHVs) present in North American *Lynx canadensis* (lynx), we obtained samples of lymphocyte-rich lynx tissues that may be sites of GHV latent infection. These samples included spleen, jejunum, lung, and bone marrow tissue from 5 *Lynx canadensis* (lynx) from Maine, USA. We confirmed that all DNA samples isolated from these tissues had amplifiable, intact DNA by PCR amplification of the glyceraldehyde-3-phosphate dehydrogenase (GAPDH) gene (data not shown). We used several pan-GHV degenerate PCR protocols for detection of GHV sequences. In our initial screen, we utilized primers that target the GHV glycoprotein B (gB) gene (Ehlers et al., 2008, 2007) and 1 of the 13 tissue samples was positive (Table 2). BLAST analysis revealed that this 452 bp GHV gB sequence from the spleen of Maine lynx 17-4019 was distinct from other gB sequences published in GenBank. The most similar sequence was the gB gene from *Lynx rufus* GHV (LruGHV1) of bobcats, with a nucleotide identity of 96%. The presence of this unique GHV gB sequence in the lynx samples provided evidence for the existence of a novel lynx GHV. We gave this putative virus the provisional name *Lynx canadensis* gammaherpesvirus 1 (LcaGHV1), in accordance with established herpesvirus naming conventions (King et al., 2012).

**Table 2.** Gammaherpesvirus sequences detected in Canada lynx by degenerate pan-GHV PCR screening of different tissues.

| **Degenerate PCR Protocol** | **Tissue Type** |  |  |  |
| --- | --- | --- | --- | --- |
|  | Spleen | Jejunum | Lung | Bone Marrow |
| Pan-GHV gB | 1/5 (17-4019<sup>a</sup>) | 0/5 | 0/3 | - |
| Pan-HV DNAPol | 0/5 | 0/5 | 0/3 | 0/3 |
| Percavirus DNAPol | 1/5 (17-4019<sup>a</sup>) | 1/5 (17-4019<sup>a</sup>) | 0/3 | 0/3 |

<sup>a</sup> Refers to identification of lynx that the tissue was isolated from

To confirm the existence of a novel GHV in lynx, we performed additional degenerate PCRs targeting the DNA polymerase (DNApol) gene. Pan-herpesviral primers commonly used for virus identification (Rose et al., 1997) were unsuccessful at amplifying DNApol sequence from 16 lynx samples (Table 2). This result was not unexpected, as we had previously found that this primer set was unable to amplify the DNApol gene of some felid GHVs (Troyer et al., 2014). With that in mind, we utilized degenerate DNApol primers designed in our previous study (Troyer et al., 2014) that are optimized for amplification of percaviruses, including GHVs of bobcats, domestic cats, and ocelots (Lozano et al., 2015). Using these primers, we found 2 of the 16 tissue samples (spleen and jejunum) to be PCR-positive, with both positives derived from the same lynx (17-4019) that was positive by gB PCR (Table 2). Sequencing and BLAST analysis revealed that the two 172 bp DNApol sequences were identical to each other and distinct from sequences in GenBank. Similar to the results for gB, the most closely related sequence was the DNApol gene of LruGHV1, with a nucleotide identity of 93%. Considering that these novel GHV sequences were isolated from the same lynx, we suspected that both sequences were from LcaGHV1.

To confirm that these gB and DNApol sequences were derived from the same virus and to obtain more sequence data for phylogenetic analysis, we designed LcaGHV1-specific primers using the gB and DNApol gene sequences and performed long-distance PCR to connect the two genes (Fig. 1A). Long-distance PCR on Maine lynx 17-4019 spleen DNA resulted in a 3.4 kb amplicon, which we sequenced using a series of primers that generated overlapping reads (GenBank <u>MH520115</u>). This result confirmed that the gB and DNApol sequences obtained above were in fact amplified from the same virus. We aligned the novel LcaGHV1 sequence with sequences from the next most closely related GHVs and found that LcaGHV1 had high nucleotide identity to the other felid percaviruses (Fig. 2). The most similar GHV to LcaGHV1 was LruGHV1 of bobcats, at 92.2% identity over this 3.4 kb region. This was closely followed by FcaGHV1 of domestic cats at 91.6% and LpaGHV1 of ocelots at 91.2%. The second GHV of bobcats, LruGHV2, was less similar at 81.6% identity.

**Figure 2.**
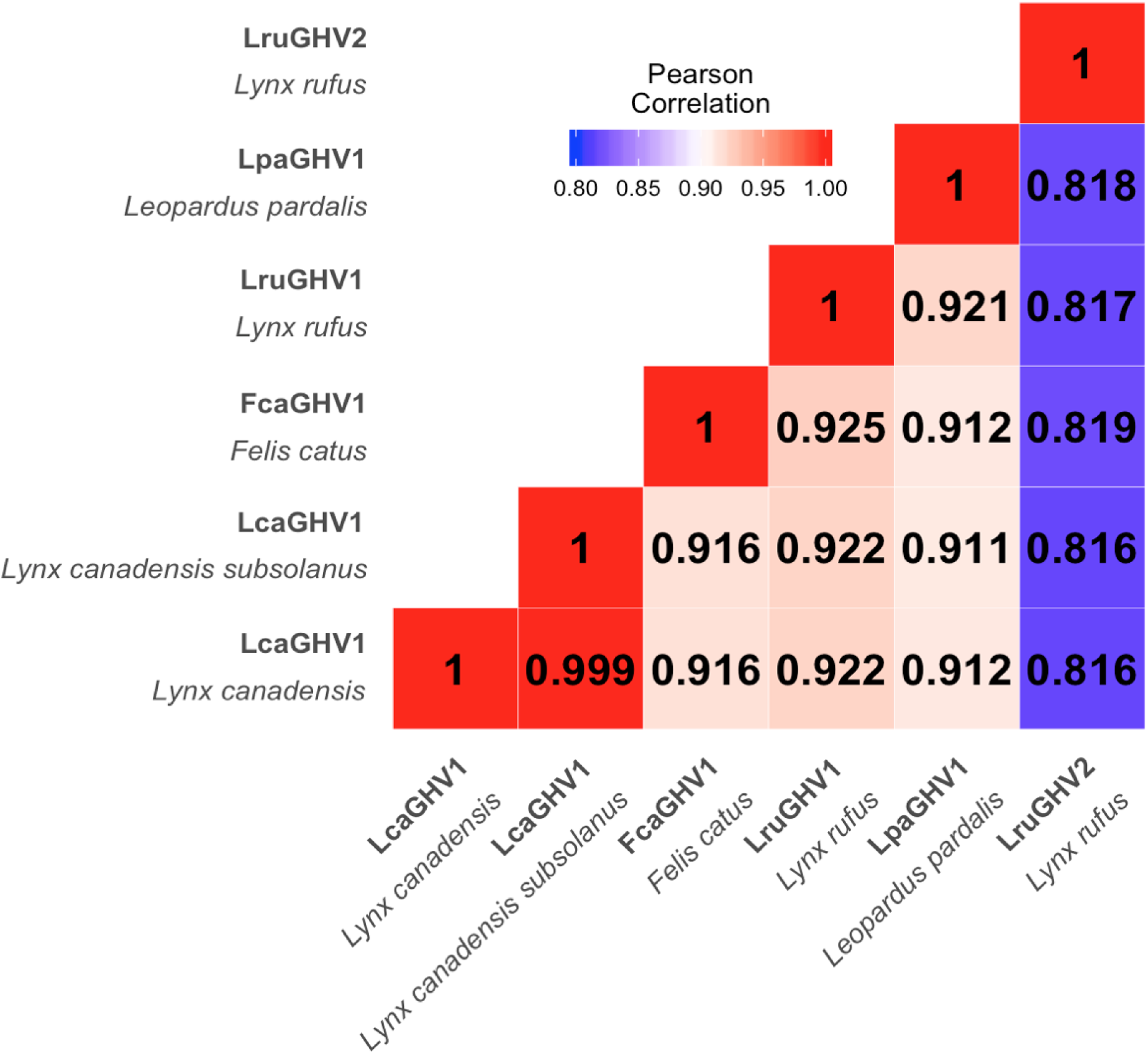
Nucleotide sequence similarity between related percaviruses with felid hosts. T-Coffee was used to align the 3.4 kb sequence spanning from gB to DNApol (as described in Fig. 1) for each virus. An identity matrix was calculated using BioEdit (Hall, 1999) and used to create this heat map.

### Prevalence of LcaGHV1 in lynx from Maine and Newfoundland

The degenerate pan-GHV primers used for identification of LcaGHV1 in this study likely have low sensitivity and specificity for LcaGHV1 and were therefore not optimal for estimating prevalence of LcaGHV1 in lynx populations. In order to reliably detect LcaGHV1 in lynx DNA samples, we designed and optimized a nested PCR protocol targeting the gB gene for sensitive detection of LcaGHV1. For this analysis, we tested lynx spleen samples from two locations; 33 *Lynx canadensis* from Maine and 41 *Lynx canadensis subsolanus* from Newfoundland, Canada (Fig. 3). Prior to LcaGHV1 detection, we confirmed that all the isolated DNA samples from both populations contained amplifiable, intact DNA by GAPDH PCR (data not shown). In total, 20 out of 74 spleen samples were LcaGHV1 PCR-positive: 13/33 from Maine and 7/41 from Newfoundland. To verify whether the amplified PCR products were derived from LcaGHV1, we sequenced the 229 nt gB amplicons for all PCR positive samples. Out of 20 positive samples, 19 had identical gB sequence to LcaGHV1 (Maine lynx 17-4019), indicating that nearly all PCR-positive samples in both Maine and Newfoundland represent the novel lynx GHV, LcaGHV1. However, the gB sequence from one Maine lynx (17-2938) was distinct from LcaGHV1 at 7 nucleotide sites and instead was identical to the sequence of LruGHV1 from bobcats (KF840716). To verify that this lynx was indeed infected with LruGHV1, which is common in bobcats and occasionally found in pumas (Troyer et al., 2014), we used degenerate PCR primers to amplify and sequence 172 nt of the GHV DNA polymerase gene. This sequence differed from LcaGHV1 at 11 nucleotide sites and only differed from the published LruGHV1 sequence at 3 sites, which is likely due to regional LruGHV1 variation (Loisel et al., 2018; Troyer et al., 2014). The identification of LruGHV1 sequence in lynx 17-2938 was not due to cross-contamination or sample switching with a bobcat as there were no bobcats examined in the animal pathology facility during the time period in which the lynx were collected and necropsied.

**Figure 3.**
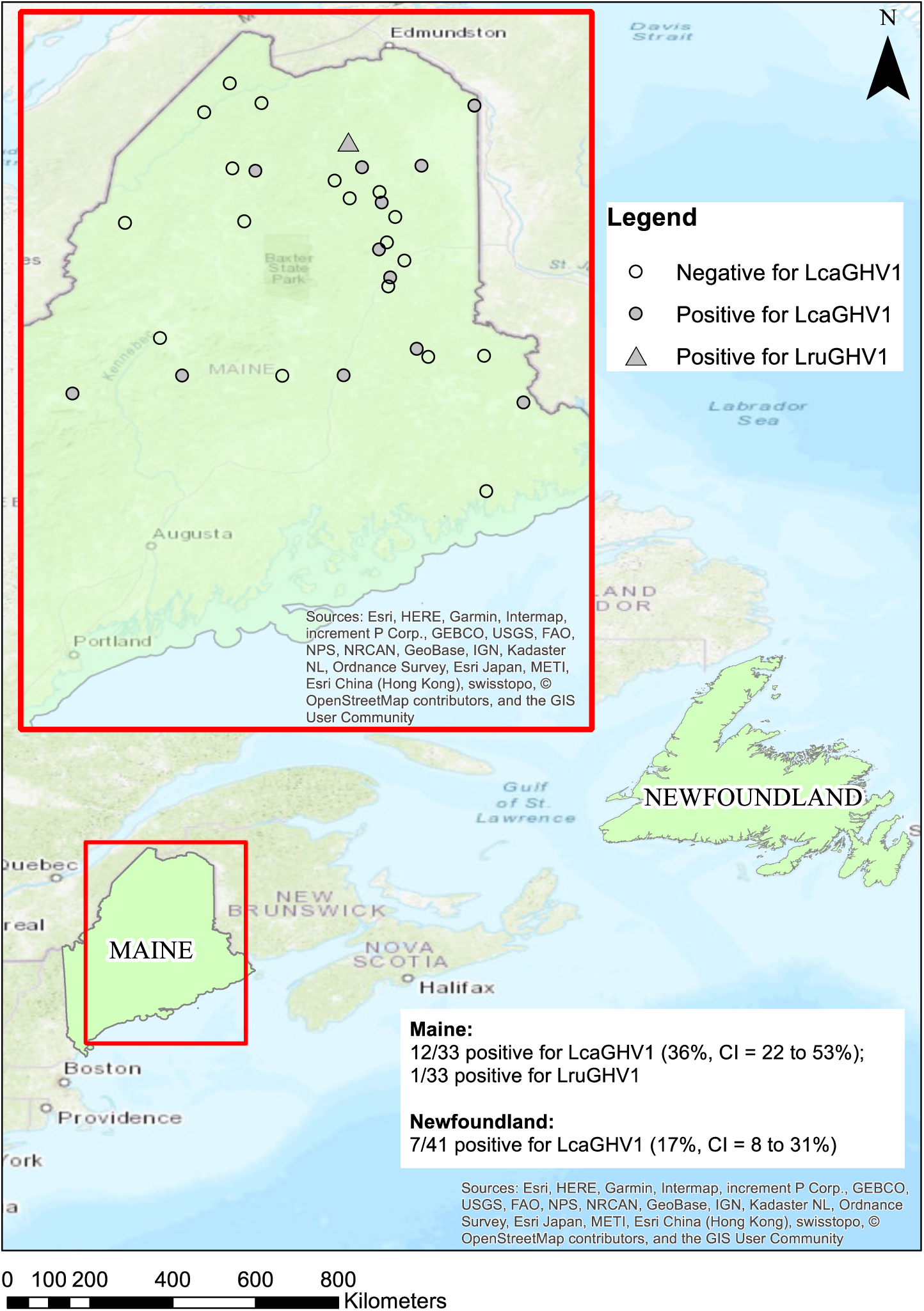
Geographic distribution of lynx samples. Both study sites are highlighted in green. For Maine the individual locations are shown for each lynx sample.

We found that 7 of 41 (17%, CI = 8-31%) Newfoundland lynx and 12 of 33 (36%, CI = 22-53%) Maine lynx were positive for LcaGHV1 (Fig. 3). One Maine lynx was found to be infected with a GHV that was clearly distinct from LcaGHV1 and nearly identical to LruGHV1 from bobcats. GPS coordinates were available for the Maine lynx samples, allowing us to plot locations of LcaGHV1 positive and negative lynx (Fig. 3). The LcaGHV1 positive samples appear relatively evenly distributed throughout the study region with no evidence of geographic clustering.

### Sequence comparison of LcaGHV1 from Maine and Newfoundland

Physical and genetic isolation of the Newfoundland island lynx subspecies *Lynx canadensis subsolanus* from mainland lynx raises the question of whether LcaGHV1 found in the Newfoundland subspecies is genetically distinct from LcaGHV1 found in mainland lynx. Short gB sequences (∼200 nt) from the 19 LcaGHV1 positive lynx in Maine and Newfoundland were all identical to each other, suggesting that LcaGHV1 from these two populations does not differ over one short region. To investigate potential GHV divergence between the two lynx populations at greater resolution, we amplified and sequenced the 3.4 kb gB to DNA polymerase region for a representative Newfoundland lynx LcaGHV1 isolate (Ly6, GenBank <u>MH520116</u>) for comparison to the same region of Maine lynx isolate 17-2019. The sequences obtained from Maine and Newfoundland differed by only 2 synonymous nucleotide substitutions (>99.9% identity, Fig. 2), suggesting that LcaGHV1 from Newfoundland lynx has not substantially diverged from LcaGHV1 in mainland lynx.

### Phylogenetic characterization of LcaGHV1

To determine the relationship of LcaGHV1 to other GHVs, we performed phylogenetic analysis. We constructed a maximum likelihood phylogenetic tree based on aligned amino acid sequences from the gB and DNApol genes of 30 previously identified GHVs, plus LcaGHV1 from *Lynx canadensis*. The clustering of LcaGHV1 provides strong evidence that it belongs to the genus *Percavirus* (Fig. 4). Mustelid GHV1 (MusGHV1), Equine GHV2 (EGHV2), and Equine GHV5 (EGHV5) are all recognized members of the genus *Percavirus*, and by tracing back to their most recent common ancestor, we see that LcaGHV1 clusters within the limits of this genus with strong bootstrap support. We had previously suggested the presence of a felid subclade within the genus *Percavirus* (Lozano et al., 2015). This analysis provides further support for this felid subclade, in that LruGHV1, FcaGHV1, LcaGHV1 and LpaGHV1 cluster together with strong bootstrap support. This tree also provides further evidence for the presence of a carnivore-host subclade within the genus *Percavirus*, with hosts from species within the order Carnivora including felids, seals, and badgers (Troyer et al., 2014).

**Figure 4.**
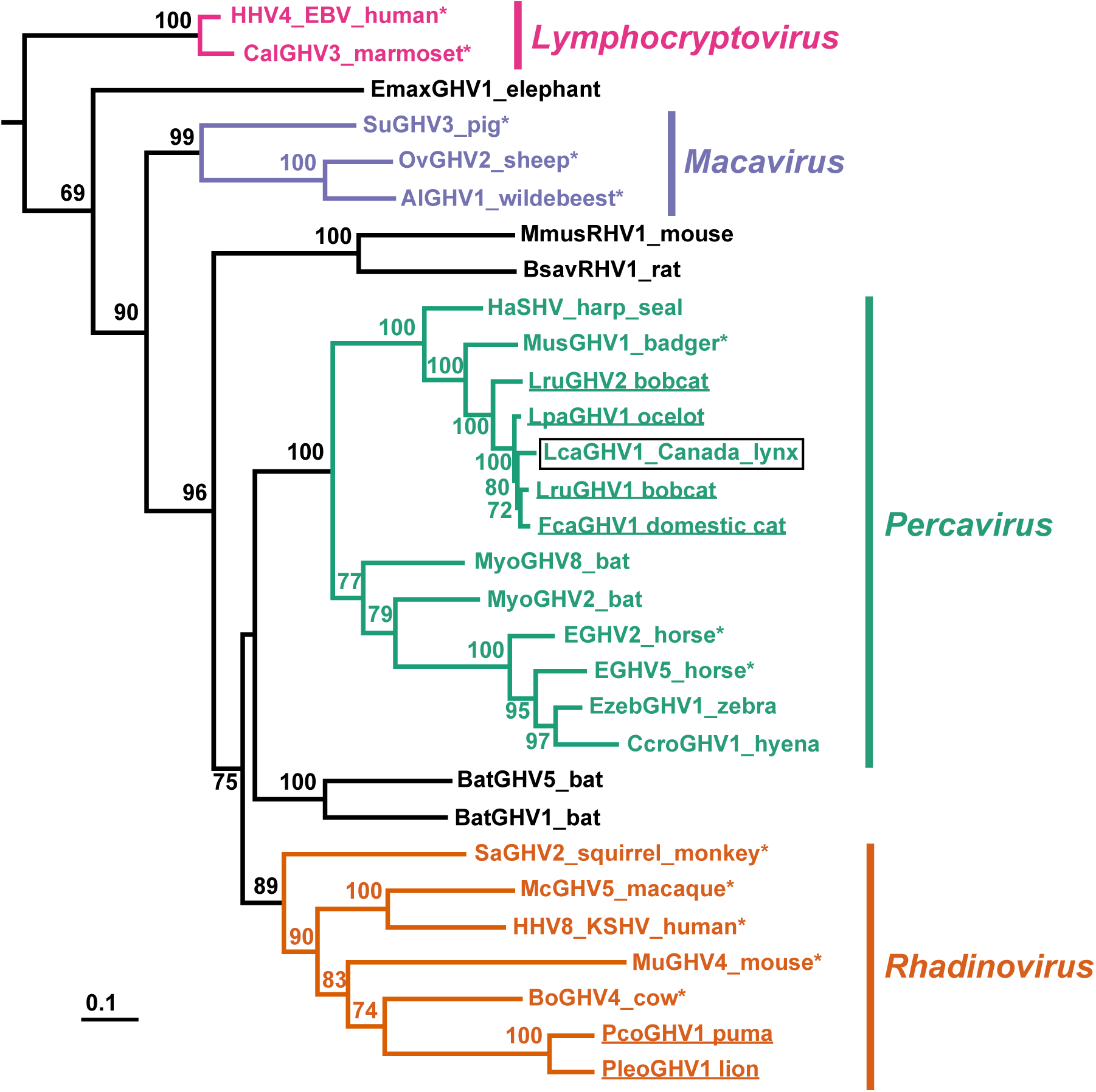
Phylogenetic analysis of various GHVs using concatenated gB and DNApol amino acid alignments. The betaherpesvirus human cytomegalovirus (human herpesvirus 5; GenBank accession no. NC006273) was used to root the tree. Bootstrap analyses were performed with 100 iterations, and support for each node is displayed (values less than 60 are not shown). Scale bar indicates genetic divergence (amino acid substitutions per site). The 4 GHV genera are labelled and were determined using the last common ancestor of accepted members of each genus (indicated with an asterisk), per the International Committee on Taxonomy of Viruses (ICTV) 2017. LcaGHV1, identified in this study, is identified with a black box. Felid-hosted GHVs are underlined. Virus abbreviations, definitions, and GenBank accession no. are as follows: HHV4, human herpesvirus 4 (Epstein-Barr virus [EBV]), NC007605; CalGHV3, callitrichine gammaherpesvirus 3, NC004367; EmaxGHV1, *Elephas maximus* gammaherpesvirus 1, EU085379; SuGHV3, suid gammaherpesvirus 3, AF478169; OvHV2, ovine herpesvirus 2, NC007646; AlGHV1, alcelaphine gammaherpesvirus 1, NC002531; BsavRHV1, *Bandicota savilei* rhadinovirus 1, DQ821581; MmusRHV1, *Mus musculus* rhadinovirus 1, AY854167; HaSHV, Harp seal herpesvirus, KF466473; MusHV1, mustelid herpesvirus 1, AF376034; LruGHV2, *Lynx rufus* GHV 2, KP721221; LpaGHV1, *Leopardus pardalis* GHV 1, KP721220; FcaGHV1, *Felis catus* GHV 1, KF840715; LruGHV1, *Lynx rufus* GHV 1, KF840716; MyoGHV8, *Myotis* gammaherpesvirus 8, KU220026; MyoGHV2, *Myotis ricketti* gammaherpesvirus 2, JN692430; EGHV2, equid gammaherpesvirus 2, NC001650; EGHV5, equid gammaherpesvirus 5, AF050671; EzebGHV1, *Equus zebra* GHV 1, AY495965; CcroGHV1, *Crocuta crocuta* GHV 1, DQ789371; BatGHV1, bat gammaherpesvirus 1, DQ788623; BatGHV5, bat gammaherpesvirus 5, DQ788629; SaGHV2, saimiriine gammaherpesvirus 2, NC001350; McGHV5, macacine gammaherpesvirus 5, NC003401; HHV8, human herpesvirus 8 (Kaposi’s sarcoma-associated herpesvirus [KSHV]), NC009333; MuGHV4, murid gammaherpesvirus 4, NC001826; BoGHV4, bovine gammaherpesvirus 4, NC002665; PcoGHV1, *Puma concolor* GHV 1, KF840717; PleoGHV1, *Panthera leo* GHV 1, DQ789370.

### Risk factors for LcaGHV1 infection in lynx

Using the prevalence data from our Maine lynx samples (n = 33), we investigated the probability of LcaGHV1 infection in relation to sex, weight, location, lungworms, lung inflammation, and heart and muscle inflammation to determine potential risk factors for infection. This analysis did not indicate any significant relationships (Table 3). Male sex has consistently been a strong risk factor for FcaGHV1 in domestic cats (Beatty et al., 2014; Ertl et al., 2015; Kurissio et al., 2018; McLuckie et al., 2016; Stutzman-Rodriguez et al., 2016; Troyer et al., 2014) but not for LruGHV1 in bobcats (Loisel et al., 2018; Troyer et al., 2014). The lack of a sex association in lynx suggests that the strong association between males and felid GHV infection may be specific to domestic and not wild cat species (Troyer et al., 2014).

**Table 3.** Potential risk factors for LcaGHV1 infection status in lynx

| Predictor | No. Samples |  | % Positive | P-value |
| --- | --- | --- | --- | --- |
|  | LcaGHV1-negative | LcaGHV1-positive |  |  |
| Sex<sup>a</sup> |  |  |  | 0.854 |
| Male | 11 | 7 | 39.0 |  |
| Female | 9 | 5 | 35.7 |  |
| Unknown | 1 | 0 | 0.0 |  |
| Location |  |  |  | 0.064 |
| Newfoundland | 34 | 7 | 17.1 |  |
| Maine | 21 | 12 | 36.4 |  |
| Lungworms<sup>a</sup> |  |  |  | 0.919 |
| Positive | 13 | 8 | 38.1 |  |
| Negative | 6 | 4 | 40.0 |  |
| Lung Inflammation<sup>a</sup> |  |  |  | 0.381 |
| Present | 11 | 5 | 31.3 |  |
| Absent | 8 | 7 | 46.7 |  |
| Heart and Muscle Inflammation<sup>a</sup> |  |  |  | 0.394 |
| Present | 12 | 5 | 29.4 |  |
| Absent | 9 | 7 | 43.8 |  |
| Weight<sup>a</sup> |  |  |  | 0.296 |
| Mean | 8.06 | 9.05 |  |  |
| n | 16 | 9 |  |  |

<sup>a</sup> Only lynx samples from Maine considered in this analysis

## Discussion

In this study, we identified novel gammaherpesviral sequence in various tissues of *Lynx canadensis* and *Lynx canadensis subsolanus*. We sequenced 3.4 kb of the novel GHVs genome and determined it to be most similar to the percavirus LruGHV1, with a nucleotide identity of 92.2%. A nucleotide identity of 93% over the same region has previously been sufficient to conclude that Suid herpesvirus 3 and 4 were distinct species, and thus we believe that this novel gammaherpesviral sequence could be a distinct GHV species (King et al., 2012). Furthermore, the frequency at which we found this novel GHV in lynx tissues provides strong evidence that the lynx is the natural host of this virus. We therefore assigned the provisional name *Lynx canadensis* gammaherpesvirus 1 (LcaGHV1), according to established naming conventions (King et al., 2012). We have now identified at least one GHV in all of the felid species present in Canada and the USA: domestic cats, bobcats, pumas, and lynx; plus, a GHV in ocelots of Central America (Lozano et al., 2015; Troyer et al., 2014). Phylogenetic analysis of LcaGHV1 supports its classification as a novel species, as well as placing LcaGHV1 within the genus *Percavirus*. We estimated the prevalence of LcaGHV1 to be 17% (n = 41, CI = 8-31%) in the Newfoundland lynx population, and 36% (n = 33, CI = 22 to 53%) in the Maine lynx population. We did not observe that factors such as sex, location, body score condition (BCS), inflammation, or infection with other agents increased the likelihood of LcaGHV1 infection and thus cannot conclude that there are any risk factors associated with this virus. We acknowledge that the relatively low sample size for which we have information on these characteristics may limit the power of this analysis.

We identified LcaGHV1 in two subspecies of lynx from two geographically distinct locations, Maine and Newfoundland. We found that over the 3.4 kb sequenced, there were only 2 nucleotide differences, both of which were synonymous. Lynx from Newfoundland are a recognized subspecies and are physically and genetically divergent from the mainland lynx population (Koen et al., 2015; Row et al., 2012; Van Zyll De Jong, 1975). This is likely due to lynx in Newfoundland being geographically isolated from the mainland population. We have previously found evidence of LruGHV1 genetic variability associated with bobcat geographic isolation. LruGHV1 in bobcats from the western United States (Colorado and California) differed by up to 4 nucleotides in a 453 nt region of gB from virus in eastern bobcats (Florida and Vermont) (Loisel et al., 2018; Troyer et al., 2014). Thus, the high similarity between LcaGHV1 isolates from each surveyed location, coupled with the fact that the two host subspecies show evidence of restricted gene flow (Koen et al., 2015; Row et al., 2012), implies that there may be an underlying mechanism by which the virus has remained relatively unchanged between these distinct locations or viral exchanges between the two populations have occurred. A low level of gene flow between Newfoundland and mainland lynx, proposed to occur by lynx crossing ice-bridges (Koen et al., 2015), suggests that viral exchange between these populations may occur through periodic contact, potentially contributing to a lack of viral diversity. Future study of lynx and LcaGHV1 genetics throughout the range of lynx in North America may provide further context for understanding the lack of LcaGHV1 genetic diversity between these two lynx sub-populations.

LcaGHV1 infection prevalence was found to be higher in the Maine population (representing the mainland population) than in the Newfoundland population (Fig. 3). There are several possible explanations for this seemingly large difference. Higher population densities have been associated with higher virus prevalence, due to increased contact and fighting (Gehrt et al., 2010; Troyer et al., 2014). Although lynx were listed as a threatened species in the conterminous United States in 2000 (Clark, 2000), improving habitat in Maine has led to rising lynx populations, with record high numbers observed in 2006 (Vashon et al., 2008). This record high population may explain the higher prevalence of LcaGHV1 in Maine. Less population information exists for lynx in Newfoundland. The sample sizes for both populations are relatively small compared to those used in previous studies, particularly for the Maine population. Expanded study of these populations is necessary to verify this potential difference.

We identified one lynx infected with LruGHV1, a virus which is typically found at high prevalence in bobcats (*Lynx rufus*) (Loisel et al., 2018; Troyer et al., 2014). While it is difficult to draw conclusions from a single animal, this suggests cross-species GHV transmission from bobcat to lynx. Bobcat to lynx transmission is plausible since bobcats are known to inhabit the area of Maine where this lynx was sampled, these species are known to interact through fighting and occasional mating (Homyack et al., 2008; Peers et al., 2013), and bobcats in nearby Vermont are infected with LruGHV1 (Loisel et al., 2018). Furthermore, we previously detected LruGHV1 DNA in blood samples from pumas (*Puma concolor*) (Troyer et al., 2014), suggesting that LruGHV1 can infect felid species in addition to bobcats. Bobcats are presumed to be the primary host species for this virus based on the high prevalence of LruGHV1 DNA in blood and spleen samples from bobcat populations in California, Colorado, Florida, and Vermont (ranging from 25% to 76% positive) (Loisel et al., 2018; Troyer et al., 2014). These findings as well as the recent finding that FcaGHV1 of domestic cats can infect Tsushima leopard cats in Japan (Makundi et al., 2018) further support the idea that felid percaviruses may naturally cross-infect multiple felid species through interspecific felid species interactions.

In this study, we designed and utilized LcaGHV1-specific nested PCR to determine prevalence in each lynx population, however this method is known to potentially underestimate the true prevalence. It is known that seropositive cats may give a negative PCR result (Stutzman-Rodriguez et al., 2016). The fact that herpesviruses establish latent infections means that there can be vanishingly small amounts of viral DNA present in the host (Barton et al., 2011; Speck and Ganem, 2010). It is therefore possible that the methods used in this study may have resulted in false negatives and underestimated the true prevalence of LcaGHV1. It is also possible that viral DNA might be present at higher levels in tissues other than spleen. Our choice to focus on spleen was based on previous studies demonstrating high levels of felid GHV DNA present in spleen compared to other tissues and blood (Beatty et al., 2014; Lozano et al., 2015). Ultimately, the prevalence results found in this study are in agreement with our previous results for prevalence of felid GHVs and provide a relatively strong estimate for a previously unknown value.

We have now identified GHVs in each felid species in northern North America. However, not all of them are percaviruses. PcoGHV1, identified in *Puma concolor*, clusters phylogenetically with rhadinoviruses (Lozano et al., 2015; Troyer et al., 2014) as confirmed in this study (Fig. 5). Thus, there are at least two distinct lineages of feline GHVs (Troyer et al., 2014). The most closely related GHV to PcoGHV1 is PleoGHV1, which infects lions (*Panthera leo*). The geographical differences between the host range of lions and pumas likely indicates that these viruses were transmitted to their hosts quite long ago, and from a potentially non-felid host (Troyer et al., 2014). In contrast, the close phylogenetic clustering of the felid percaviruses, and similarities in geographical distribution of felid hosts throughout North America, could indicate more recent cross-species transmission events. Alternatively, the branching pattern seen in the evolution of felids is nearly identical to that of their respective GHVs (Johnson et al., 2006), suggesting speciation could account for the observed felid percavirus subclade. More recently, FcaGHV1 has been identified in Europe (Ertl et al., 2015), Japan (Tateno et al., 2017), and Brazil (Kurissio et al., 2018), suggesting world-wide distribution. Though it is not clear whether this world-wide distribution is natural or whether it is due to human activity, especially since domestic cats are pets in many parts of the world. Since there is less human-influenced movement of wild felids such as Canada lynx and Eurasian lynx (*Lynx lynx*), we believe that further insights into the evolution of feline GHVs could be uncovered by screening additional felid species such as the Eurasian lynx for novel GHVs. Given the identification of three distinct percaviruses in North American species of the *Lynx* genus (LruGHV1, LruGHV2, and now LcaGHV1), we would expect to find a percavirus in Eurasian lynx as well.

We did not find any statistically significant risk factors for LcaGHV1 infection (Table 3). We and others had previously found that being male and of older age were significant risk factors for FcaGHV1-positive status in domestic cats (Beatty et al., 2014; Troyer et al., 2014), and this relationship has been confirmed by other studies (Ertl et al., 2015; Kurissio et al., 2018; McLuckie et al., 2016; Stutzman-Rodriguez et al., 2016). This relationship between sex and infection status was not observed in wild felids (pumas and bobcats), however the association with higher age was (Troyer et al., 2014). Similarly, in the present study of wild lynx, we found no association between sex and GHV infection status. We could not evaluate animal age as a risk factor since this information was only available for a few lynx. Beyond the risk factors that we had previously examined, we also wanted to investigate the relationship between LcaGHV1 infection and lung inflammation. While the pathogenesis of other percaviruses is largely unknown, equine GHVs (EGHV2 and EGHV5) have been associated with respiratory disease (Fortier et al., 2010). We did not find any significant relationship between being LcaGHV1-positive and lung inflammation (Table 3). It is also worth noting that while EGHV2 and EGHV5 are known to have tissue tropism in the lungs (Fortier et al., 2010), we did not detect LcaGHV1 in any of the lynx lung tissue sampled (Table 2). Finally, we did not find any significant relationship between infection status and lynx health (using body condition score and weight as proxies for lynx health) (Table 3). This suggests that LcaGHV1 may not cause an overt disease phenotype. However, we have to be cautious about making strong conclusions as the number of samples for statistical analysis was relatively small. The results do not rule out LcaGHV1 from causing disease in lynx and further characterization of risk factors and pathogenesis is warranted.

In conclusion, we have identified a novel felid percavirus that commonly infects lynx in northeastern North America. This work contributes to our understanding of GHV evolutionary history and may have future implications for the health and ecology of lynx and other felids in North America. Future studies into risk factors and pathogenesis of GHVs in North American felids will be necessary to determine whether the health of these animals is at risk.

## Acknowledgements

This work was supported by an operating grant from the Natural Sciences and Engineering Research Council of Canada (NSERC) to RMT and by infrastructure funds from the University of Western Ontario. Lynx carcasses from Maine were provided by the Maine Department of Inland Fisheries and Wildlife with funding provided by Federal Aid in Wildlife Restoration, Maine State Wildlife Grants, Maine Wildlife Conservation and Restoration Program, and Maine Section 6 Endangered Species Grants.

